# Comparative aerosol and surface stability of SARS-CoV-2 Variants of Concern

**DOI:** 10.1101/2022.11.21.517352

**Authors:** Trenton Bushmaker, Claude Kwe Yinda, Dylan H. Morris, Myndi G. Holbrook, Amandine Gamble, Danielle Adney, Cara Bushmaker, Neeltje van Doremalen, Robert J. Fischer, Raina K. Plowright, James O. Lloyd-Smith, Vincent J. Munster

**Affiliations:** Laboratory of Virology, Division of Intramural Research, National Institute of Allergy and Infectious Diseases, National Institutes of Health, Hamilton, MT, USA; Dept. of Microbiology and Cell Biology, Montana State University, Bozeman, MT, USA; Dept. of Ecology and Evolutionary Biology, University of California, Los Angeles, Los Angeles, CA, USA; Fogarty International Center, National Institutes of Health, Bethesda, MD, USA; Bitterroot Health - Daly Hospital, Hamilton, MT, USA

## Abstract

SARS-CoV-2 is transmitted principally via air; contact and fomite transmission may also occur. Variants-of-concern (VOCs) are more transmissible than ancestral SARS-CoV-2. We find that early VOCs show greater aerosol and surface stability than the early WA1 strain, but Delta and Omicron do not. Stability changes do not explain increased transmissibility.

---

Since the initial emergence of SARS-CoV-2 (Lineage A), new lineages and variants have emerged, typically replacing previously circulating lineages (1). Five virus variants had been characterized as Variants of Concern (VOCs) by the World Health Organization (2, 3). To assess whether the transmission advantage of new VOCs may have arisen partly from changes in aerosol and surface stability, we directly compared the Lineage A ancestral virus (WA1 isolate) with VOCs from later timepoints during the pandemic.

## The Study

We evaluated the stability of SARS-CoV-2 variants in aerosols and on and on high density polyethylene and estimated their decay rates using a Bayesian regression model (see the Methods section in the Supplementary Appendix). Aerosols (<5 μm) containing SARS-CoV-2 variants of 10^5.75^ - 10^6^ TCID_50_ (50% tissue-culture infectious dose [TCID_50_] per milliliter [mL]) were generated with a three-jet Collison nebulizer and fed into a Goldberg drum to create an aerosolized environment (supplemental video). On polyethylene, 50 μL of a solution containing roughly 10^5^ TCID_50_ of virus was applied to measure surface stability.

For aerosol stability, we directly compared the exponential decay rate of different SARS-CoV-2 isolates (Table 1) at time points 0, 3 and 8 hours; the 8-hour time point was chosen to maximize information on decay rate given the observed 3-hour decay. Experiments were performed as single runs (0-to-3 or 0-to-8-hours), with sample collection at the start and end points, to minimize virus loss and humidity changes from repeat sampling. All runs were conducted in triplicate. To estimate quantities of sampled virus, air samples collected at 0, 3 or 8 hours post-aerosolization were analyzed by reverse transcription polymerase chain reaction (qRT-PCR) for the SARS-CoV-2 E-gene to determine the amount of genome copies within the samples. To determine the remaining concentration of viable (infectious) SARS-CoV-2 virions, the samples were titrated on Vero E6 cells. Exponential decay of infectious virus was estimated relative to the amount of genome copies to account for particle settling or other physical loss of viruses.

**Table 1.**
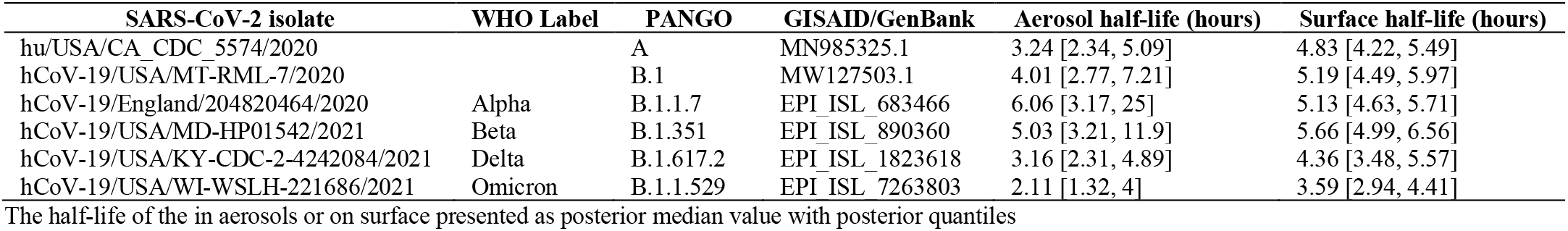
SARS-CoV-2 isolates used in this study with the observed aerosol and surface halflives.

| SARS-CoV-2 isolate | WHO Label | PANGO | GISAI/GenBank | Aerosol half-life (hours) | Surface half-life (hours) |
| --- | --- | --- | --- | --- | --- |
| hu/USA/CA_CDC_5574/2020 |  | A | MN985325.1 | 3.24 [2.34, 5.09] | 4.83 [4.22, 5.49] |
| hCoV-19/USA/MT-RML-7/2020 |  | B.1 | MW127503.1 | 4.01 [2.77, 7.21] | 5.19 [4.49, 5.97] |
| hCoV-19/England/204820464/2020 | Alpha | B.1.1.7 | EPI_ISL_683466 | 6.06 [3.17, 25] | 5.13 [4.63, 5.71] |
| hCoV-19/USA/MD-HP01542/2021 | Beta | B.1.351 | EPI_ISL_890360 | 5.03 [3.21, 11.9] | 5.66 [4.99, 6.56] |
| hCoV-19/USA/KY-CDC-2-4242084/2021 | Delta | B.1.617.2 | EPI_ISL_1823618 | 3.16 [2.31, 4.89] | 4.36 [3.48, 5.57] |
| hCoV-19/USA/WI-WSLH-221686/2021 | Omicron | B.1.1.529 | EPI_ISL_7263803 | 2.11 [1.32, 4] | 3.59 [2.94, 4.41] |

The half-life of the in aerosols or on surface presented as posterior median value with posterior quantiles

We were able to recover viable SARS-CoV-2 virus from the drum for all VOCs (Figure 1A). The quantity of viable virus decayed exponentially over time (hence a linear decrease in the log_10_TCID_50_ per liter of air over time (Figure 1B)). The half-life of the ancestral lineage WA1 in aerosols (posterior median value [2.5% to 97.5% posterior quantiles]) was 3.24 [2.34, 5.09] hours. The B.1, Alpha and Beta viruses appeared to have longer half-lives than WA1: 4.01 [2.77, 7.21] hours for B.1, 6.06 [3.17, 25] hours for Alpha, and 5.03 [3.21, 11.9] hours for Beta. The Delta variant displayed a half-life similar to that of WA1: 3.16 [2.31, 4.89] hours. The Omicron variant displayed a similar or decreased half-life compared to WA1: 2.11 [1.32, 4] hours (Figure 1B). To better quantify the magnitude and certainty of the change, we computed the posterior of the ratio for variant half-life to WA1 half-life for each variant (Figure 1C). Estimated ratios were 1.24 [0.697, 2.42] for B.1, 1.88 [0.846, 7.64] for Alpha, 1.56 [0.842, 3.99] for Beta, 0.981 [0.558, 1.69] for Delta, and 0.652 [0.344, 1.33] for Omicron. That is, initial spike protein divergence from WA1 (heuristically quantified by the number of non-synonymous amino acid substitutions) appeared to produce increased relative stability, but further evolutionary divergence reverted stability back to that of WA1, or even below it (Figure 1C, Figure S1, Table S1).

**Figure 1.**
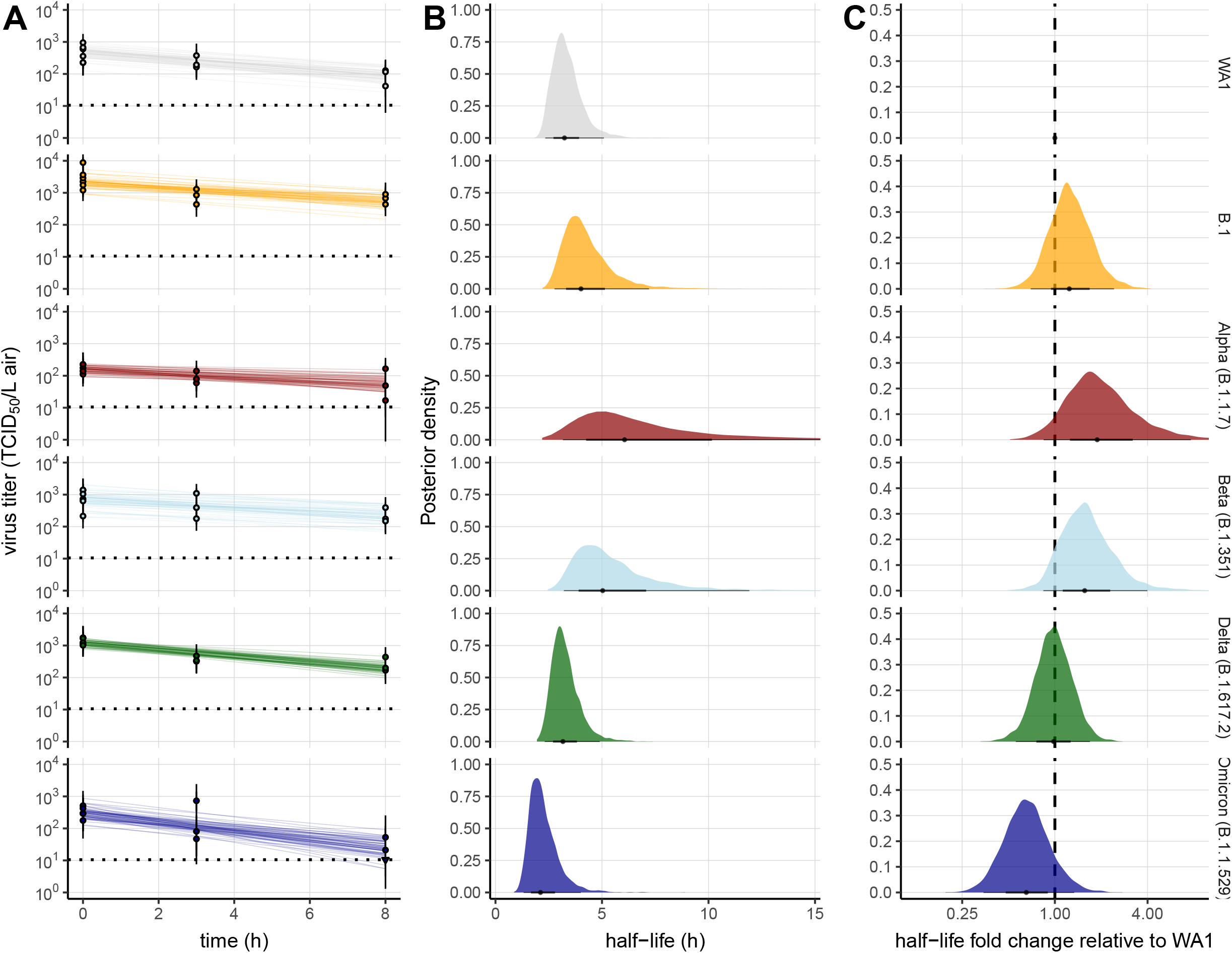
SARS-CoV-2 variant exponential decay in aerosolized form and corresponding halflives. A: Regression lines showing predicted decay of log_10_ virus titer over time compared to measured (directly inferred) virus titers. Points show posterior median measured titers; black lines show a 95% credible interval. Point at 3h and 8h are shifted upward by the estimated physical / non-inactivation loss of virus, as estimated from qPCR data (see Supplementary text Methods) to enable visual comparison with the predicted exponential decay curve (which reflects inactivation only). Colored lines are random draws from the joint posterior distribution of the exponential decay rate (negative of the slope) and intercept (initial virus titer); this visualizes the range of possible decay patterns for each experimental condition. 10 lines plotted for each drum run (thus 60 per panel). B: Inferred virus half-lives by variant. Violin plots show the shape of the posterior distribution. Dots show the posterior median half-life estimate and black lines show a 68% (thick) and 95% (thin) credible interval. C: Inferred ratio of variant virus half-lives to that of WA1 (fold-change), plotted on a logarithmic scale and centered on 1 (no change). Dot shows the posterior median estimate and black lines show a 68% (thick) and 95% (thin) credible interval.

Next, we investigated the surface stability of VOCs compared to the ancestral variant on polyethylene. Again, all variants exhibited exponential decay, as indicated by linear decrease in the log_10_TCID_50_/mL over time (Figure 2A). We found a half-life (posterior median [2.5% and 97.5% quantiles]) of 4.83 [4.22, 5.49] hours for WA1, similar to our previous estimates (Figure 2B) (4). B.1, Alpha, and Beta had slightly longer half-lives: 5.19 [4.49, 5.97] hours for B.1, 5.13 [4.63, 5.71] hours for Alpha, and 5.66 [4.99, 6.56] hours for Beta (Figure 2B). As in aerosols, Delta had a half-life similar to WA1 of 4.36 [3.48, 5.57] hours, and Omicron had a somewhat shorter half-life 3.59 [2.94, 4.41] hours (Figure 2B). To quantify the strength and statistical discernibility of the pattern, we calculated posterior probabilities for the half-life ratio relative to WA1 (Figure 2C). B.1, Alpha, and Beta had half-life ratios to WA1 of 1.08 [0.886, 1.3], 1.06 [0.905, 1.26], and 1.18 [0.978, 1.42], respectively. Delta had a ratio of 0.902 [0.699, 1.19], and Omicron 0.744 [0.582, 0.948].

**Figure 2.**
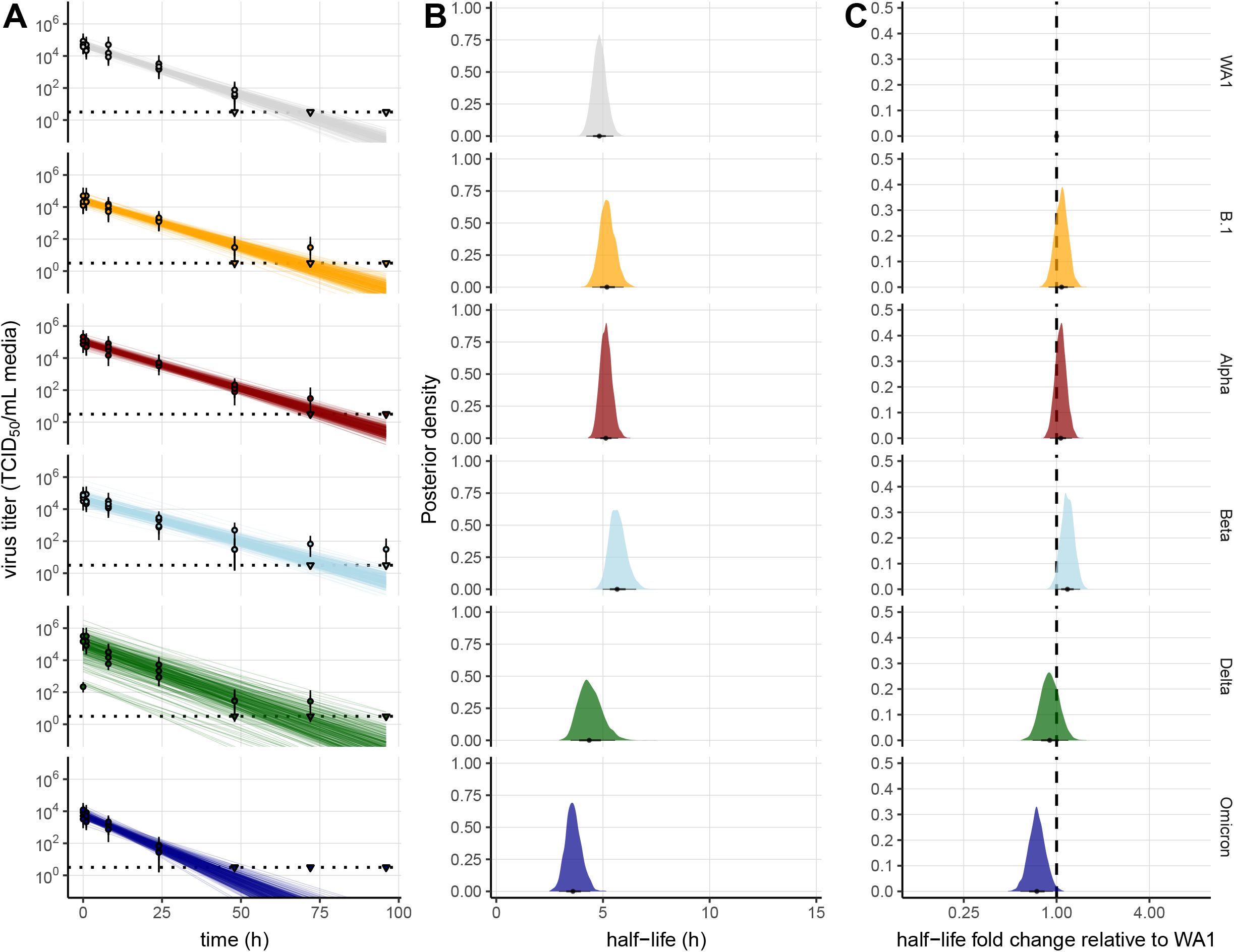
SARS-CoV-2 variant exponential decay on an inert surface and corresponding halflives. A: Regression lines showing predicted decay of log_10_ virus titer over time compared to measured (directly inferred) virus titers. Points show posterior median measured titers; black lines show a 95% credible interval. Colored lines are random draws from the joint posterior distribution of the exponential decay rate (negative of the slope) and intercept (initial virus titer); this visualizes the range of possible decay patterns for each experimental condition. 10 lines plotted per panel for each sampled titer. B: Inferred virus half-lives by variant. Violin plots show the shape of the posterior distribution. Dots show the posterior median half-life estimate and black lines show a 68% (thick) and 95% (thin) credible interval. C: Inferred ratio of variant virus half-lives to that of WA1 (fold-change), plotted on a logarithmic scale and centered on 1 (no change). Dot shows the posterior median estimate and black lines show a 68% (thick) and 95% (thin) credible interval.

## Conclusion

Several studies have analyzed the stability of SARS-CoV-2 in aerosols in a Goldberg rotating drum.(4–7) In general, these studies focused on the duration over which infectious virus could be detected. In one study, virus was detected after 16 hours (6). Studies using the VOCs B.1 and B.1.1.7 did not detect notable differences in aerosol stability compared to ancestral strains (8).

Here, we paired a model-optimized experimental design with Bayesian hierarchical analysis to systematically measure virus half-life across six SARS-CoV-2 variants and directly estimate relative half-lives with full error propagation. We found a small initial increase in aerosol stability from ancestral WA1 to the B.1, Alpha and Beta variants, with some statistical uncertainty. However, we found that Delta has a half-life similar to WA1, and Omicron a shorter one. In surface measurements, the VOCs followed the same pattern of relative stability, confirming that the overall stability of SARS-CoV-2 variants is determined by similar factors in aerosols and on surfaces.(9)

Our study suggests that aerosol stability is likely not a major factor driving the increase in transmissibility observed with several VOCs.(10, 11) The early rise in stability for B.1 and its descendants Alpha and Beta may have been incidentally caused by selection for other viral traits that favored higher transmission. Epidemiological and experimental studies suggest that the window for transmission is typically relatively short (under an hour), and that a modest change in aerosol half-life will not have a discernible impact on the epidemiological level (12). However, in specific contexts of enclosed spaces, it will remain important to understand the temporal profile of transmission risks after the release of aerosols containing SARS-CoV-2 from an infected individual.

Whereas evolutionary selection for prior variants favored high transmission to and from naïve humans(13), since late 2021 global population-level selection have favored antigenic change, and the consequent ability to transmit to and from non-naïve individuals (14, 15). But as the example shows, either increased transmissibility in naïves or adaptive antigenic evolution may come at a tolerable cost in environmental stability. Overall, the minor differences in the environmental stability between isolates of different VOCs in aerosols or on surfaces are unlikely to be driving variant population-level epidemiology.

## Acknowledgements

We would like to thank Michele Adams for clinical specimen acquisition, Friederike Feldmann for experimental support, and Austin Athman for creation of the video. The following SARS-CoV-2 isolates were obtained through: CDC: SARS-CoV-2/human/USA/WA-CDC-WA1/2020, Lineage A. BEI Resources, NIAID, NIH: SARS-CoV-2 variant Alpha (B.1.1.7) (hCoV320 19/England/204820464/2020, EPI_ISL_683466), contributed by Bassam Hallis, and variant Delta (B.1.617.2/) (hCoV-19/USA/KY-CDC-2-4242084/2021, EPI_ISL_1823618). Variant Beta (B.1.351) isolate name: hCoV-19/USA/MD-HP01542/2021, EPI_ISL_890360, was contributed by Johns Hopkins Bloomberg School of Public Health: Andrew Pekosz. Variant Omicron (B.1.1.529. BA.1) isolate name: hCoV-19/USA/GA-EHC-2811C/2021, EPI_ISL_7171744, was contributed Emory University, Emory Vaccine Center: Mehul Suthar. We thank Andrew Pekosz, Mehul Suthar, Emmie de Wit, Brandi Williamson, Sujatha Rashid, Bassam Hallis, Ranjan Mukul, Kimberly Stemple, Bin Zhou, Natalie Thornburg, Sue Tong, Stacey Ricklefs, Sarah Anzick for gracefully sharing viruses or propagating and sequence confirming virus stocks. This research was supported by the Intramural Research Program of the National Institute of Allergy and Infectious Diseases (NIAID), National Institutes of Health (NIH) and Defense Advanced Research Projects Agency DARPA PREEMPT # D18AC00031. Contributions of D.H.M., A.G. and J.L.S. were further supported by the National Science Foundation (DEB-1557022) and the UCLA AIDS Institute and Charity Treks. This work was part of NIAID’s SARS-CoV-2 Assessment of Viral Evolution (SAVE) Program.

Mr. Bushmaker is a biologist in NIAIDs Laboratory of Virology and is interested in high containment pathogens. Dr. Yinda is a postdoctoral research fellow in NIAIDs Laboratory of Virology. He is interested in emerging viruses, and their transmissions potential.

## Author contributions

Conceptualization: T.B., C.K.Y., A.G., D.M., R.J.F, V.M., and J.L.S.; methodology: T.B., C.K.Y., M.H., and D.R.A., and R.J.F.; resources: C.I.B. and N.v.D.; supervision: V.J.M. and R.K.P.; data curation: T.B., C.K.Y., D.M., R.J.F., and A.G.; writing: T.B., C.K.Y., A.G., D.M., R.K.P., V.M., and J.L-S.; visualization: T.B., C.K.Y., and D.M.

## Supplementary Appendix

### Materials and Methods

#### Cells and viruses

SARS-CoV-2 strains were passaged once on VeroE6 cells maintained in DMEM supplemented with 10% FBS, 2 mM L-glutamine, 100 U/ml penicillin and 100 μg/ml streptomycin.

The ancestral WA1 (lineage A) strain hu/USA/CA_CDC_5574/2020 (MN985325.1) was provided by CDC, Atlanta, USA. The B.1 hCoV-19/USA/MT-RML-7/2020 (GISAID# EPI_ISL_591054, MW127503.1) was derived from a clinical specimen obtained from Bitterroot Health - Daly Hospital Hamilton, USA. For the VOCs, Alpha variant B.1.1.7 hCOV_19/England/204820464/2020, NR-54000 (GISAID#EPI_ISL_683466) was obtained through BEI Resources, NIAID, NIH: Severe Acute Respiratory Syndrome-Related Coronavirus 2, contributed by Dr. Bassam Hallis. Beta variant B.1.351 hCoV-19/USA/MD-HP01542/2021 (GISAID# EPI_ISL_890360) was acquired from Dr. Tulio de Oliveira and Dr. Alex Sigal at the Nelson R. Mandela School of Medicine, UKZN. The Delta variant B.1.617.2 hCoV-19/USA/KY-CDC-2-4242084/2021 (GISAID# EPI_ISL_1823618) was obtained from BEI and the Omicron variant B.1.1.529 hCoV-19/USA/WI-WSLH-221686/2021 ((GISAID# EPI_ISL_7263803 was obtained from Drs. Peter Halfmann and Yoshihiro Kawaoka at the University of Wisconsin – Madison, USA (Table 1).

All virus stocks were propagated in VeroE6 cells in DMEM supplemented with 2% fetal bovine serum (FBS), 2 mM L-glutamine, 100 U/ml penicillin and 100 μg/ml streptomycin. Stocks were harvested between Day 4 - 6, dependent of the cytopathic effect. Supernatant was collected to be centrifuged at 1200 rpm for 8 minutes at room temperature centrifugation and frozen at −80°C. To determine if virus isolates genomes were identical to those deposited in GenBank and/or GISAID, we perform deep sequencing with the Illumina MiSeq system using nano 300-cycle chemistry (Illumina). The data is presented in Table 1.

#### SARS-CoV-2 stability in aerosol - Goldberg drum exposure and sample analysis

Droplet nuclei size particles (<5 μm) were generated using a 3-jet Collison nebulizer (CH Technologies) containing 10^5.75^ - 10^6^ TCID_50_/mL in 10 mL of DMEM supplemented with 2% FBS. The inoculum fed into a rotating Goldberg drum (Biaera Technologies) to create an aerosolized environment. The drum system was prepared until a starting environment of 65% relative humidity (RH) and a temperature of 21-23°C was reached for all SARS-CoV-2 Goldberg drum runs. Aerosols were maintained in suspension with a rotation of 3 mph to overcome terminal settling velocity.

Three independent replicates were performed for each of the respective timepoint, either a 3-hour or 8-hour run for each of the SARS-CoV-2 strains assessed in this study. For each independent run, samples were collected at 0 and 3-hour or 0 and 8-hour post aerosol generation. Samples were collected by drawing air at 6 LPM for 30 secs onto a 47mm gelatin filter (Sartorius). Filters were dissolved in 10 mL of DMEM containing 10% FBS at 37°C. Samples were frozen at −80°C until assessment.

Aerosol samples were quantified using qRT-PCR as previously described [17]. In short, 140 μL of sample was utilized for RNA extraction using the QIAamp Viral RNA Kit (Qiagen) using QIAcube HT automated system (Qiagen) with an elution volume of 150 μL. SARS-CoV-2 was detected using the E gene assay in a qRT-PCR (Corman et al., 2020) using 5 ul of input RNA and the TaqMan™ Fast Virus One-Step Master Mix (Applied Biosystems) and run on a QuantStudio 6 Flex Real-Time PCR System (Applied Biosystems). 10-fold dilutions SARS-CoV-2 E gene run-off transcripts 10-fold dilutions with known genome copies were run in parallel to allow calculation of genome copies in samples. Infectious virus titers were determined by end-point titration on VeroE6 cells and TCID_50_/mL was calculated using method of Spearman-Karber on VeroE6 cells.

#### SARS-CoV-2 stability on surface

Surface stability was evaluated on 15 mm polypropylene at 21-23°C/40% RH. 50 μL of virus stock containing 105 TCID_50_/mL was deposited (7-10 drops) on the surface of a disc. At predefined time-points viable virus was recovered by rinsing with 1 mL of Dulbecco’s modified Eagle’s medium (Sigma-Aldrich, St, Louis, MO) supplemented with 2% fetal bovine serum, 1 mM L-glutamine, 50 U/ml penicillin and 50 μg/ml streptomycin (2% DMEM) and frozen at −80°C until titrated. Three replicate experiments were performed for each surface and infectious virus titers were determined by end-point titration as described above.

### Supplementary figures

**Figure S1.**
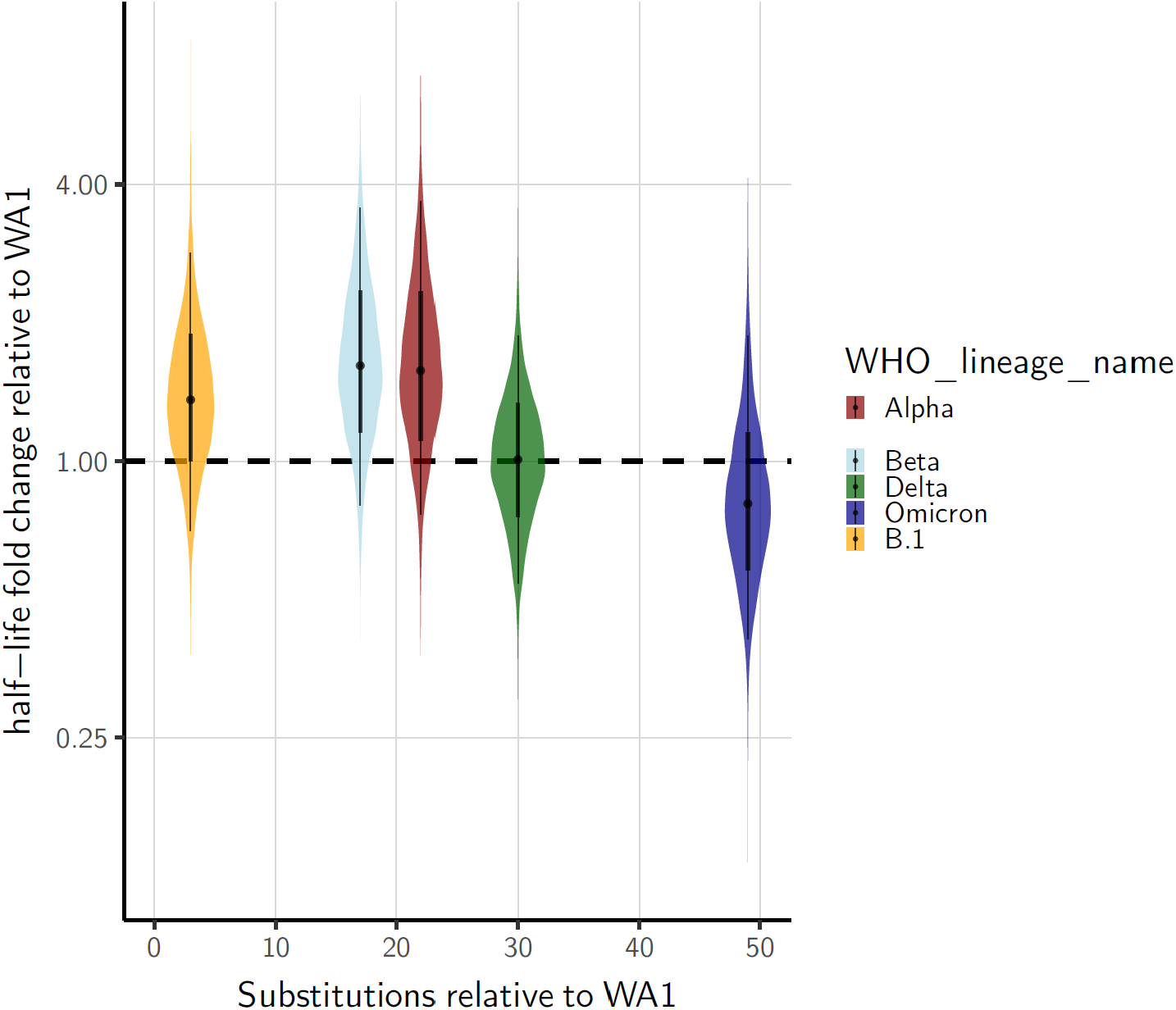
Correlation of the half-lives in proportion of the amino acid substitution in the genome of the ancestral cohort or Variants of Concern relative to the WA1 Lineage A.

**Table S1.**
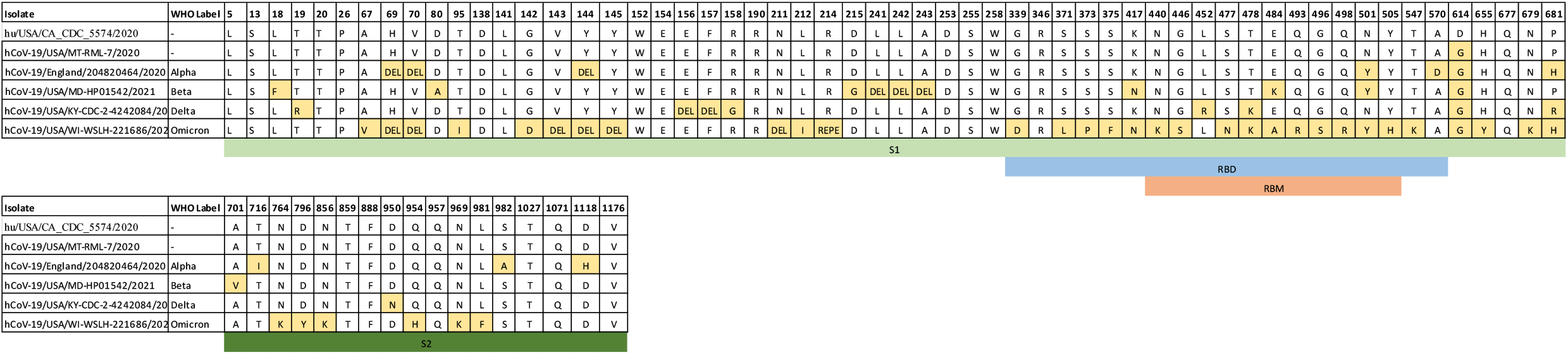
Amino acid substitutions and deletions in the S1 and S2 regions of the spike glycoprotein of the Variants of Concern relative to the WA1 Lineage A.

## Supplementary text: Bayesian infererence methods

### 1 Conceptual overview

Building on our prior work^1–3^, we inferred individual titers and virus half-lives in a Bayesian framework, modeling the positive or negative status of individual observed titration wells according to a Poisson single-hit process^4^. This can then be used either to infer individual titers or to fit an exponential decay rate (equivalent, a half-life) to a set of samples taken at different timepoints. In the latter case, we jointly infer the decay rate and the individual titers, for maximally principled error propagation. The reason we also estimate individual titer values (without any assumptions about their relationship or the decay process) is that this allows us to check goodness-of-fit of the exponential decay model.

### 2 Notation

In the text that follows, we use the following mathematical notation.

#### 2.1 Logarithms and exponentials

log(*x*) denotes the logarithm base *e* of *x* (sometimes called ln(*x*)). We explicitly refer to the logarithm base 10 of *x* as log_10_(*x*). exp(*x*) denotes *e^x^*.

#### 2.2 Probability distributions

The symbol ~ denotes that a random variable is distributed according to a given probability distribution. So for example

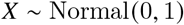

indicates that the random variable *X* is normally distributed with mean 0 and standard deviation 1.

We parameterize normal distributions as:

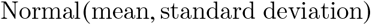

We parameterize positive-constrained normal distributions (i.e. with lower limit 0) as:

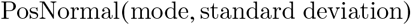

We parameterize Poisson distributions as:

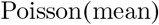

### 3 Titer inference

For both surface and aerosol samples, we estimated individual sample infectious virus titers directly from titration well data as previously described^2^, using a weakly informative Normal prior on the true virus concentration *v_i_* in units of TCID_50_/0.1mL (since well inocula were 0.1 mL):

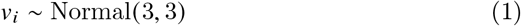

### 4 Surface half-life inference

Similarly, we inferred half-lives of infectious virus on surfaces using the method previously described in^2^, which allows us to account for variation in initial virus deposition on individual coupons, among other sources of experimental error. We used the following priors.

Log half-lives log(*h_i_*) for each experimental condition *i*:

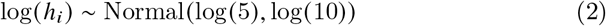

Mean initial log_10_ virus titers 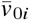 for each experimental condition *i*:

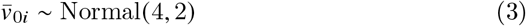

Experiment-specific standard deviations *σ_i_* of initial initial log_10_ titers *v*_0*ij*_ about the mean 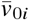 for each experimental condition *i*:

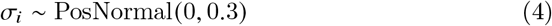

### 5 Aerosol half-life inference

To conduct aerosol half-life inference, we had to account for settling and other loss of virus unrelated to virus inactivation. We did this by incorporating qPCR measurements of virus genome quantity. For each drum run *j* of experimental condition *i,* we estimated *L_ij_*, the non-inactivation loss of infectious virus for the final (*t* = 3 h or *t* = 8 h) sample 1*ij* relative to the initial *t* = 0 h sample 0_*ij*_ (in units of log_10_ infectious virus), by the change in sample CT values *C_ij_*:

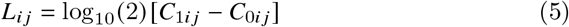

For each drum run *j*, we then predicted the measured final infectious virus titer *v*_1*ij*_ given the *t* = 0*h* measurement *v*_0*ij*_ as:

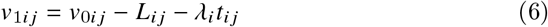

where *t_ij_* is the timepoint for the second sample (3 h or 8 h) and *λ_i_* is the exponential decay rate in log_10_ infectious virus per hour, calculated from the half-life as:

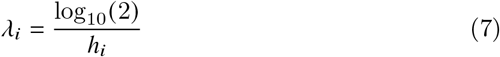

We assume that the initial sampled titers *v*_0*ij*_ for each individual drum run *j* of experiment *i* are distributed about an inferred experiment-specific mean 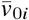, with an inferred experiment-specific standard deviation *σ_i_*:

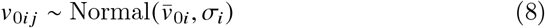

We treated observed titration wells for both *v*_0*ij*_ and *v*_1*ij*_ according to the same Poisson single-hit process previously described and used to estimate individual titers and surface half-lives.

We used the following priors.

Log half-lives log (*h_i_*) for each experimental condition *i*:

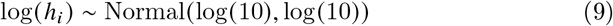

Mean initial virus titers 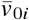 (in units of log_10_ TCID_50_/0.1 mL titrated sample):

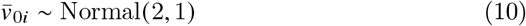

Standard deviations *σ_i_* of individual initial titers *v*_0*ij*_ about the experiment mean 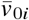:

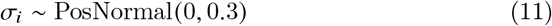

### 6 Computational methods

As previously described, we fit the models described above our data using Stan^5^, which implements a No-U-Turn Sampler^6^. We inferred all parameters jointly for all models. We ran 4 parallel Markov chains with 1000 iterations of warmup followed by 1000 sampling iterations, resulting in a total of 4000 posterior samples for each inference model. We assessed chain mixing and convergence by inspecting trace plots and confirming sufficient effective sample size and lack of divergent transitions.

We created visualizations and tables in R using `ggplot2`^7^, `ggdist`^8^, and `tidybayes`^9^.

### 7 Methodological discussion

The principal difference between the drum and the surface experiments is that in the drum experiments we directly sample *v*_0*ij*_, as this can be done non-destructively (where it cannot be done with an individual surface sample).

Note that the *t* = 0 h sample in the aerosol experiments occurs after a drum equilibriation period, and thus after any physical loss from that occurs during the aerosolization process and any rapid initial loss of infectious virus, as has been reported in other studies of aerosolized virus^10^.

Except for very near-field airborne exposure (e.g. a person shouting in another’s face), the transmission-relevant half-life of infectious virus in aerosols is the quasi-equilibrium half-life after any rapid initial loss has occurred. This later half-life as the one our experiment is designed to measure (note that our *t* = 0 h titers are much lower than our stock solution.

Similarly, it is important to note that real-world depositions in aerosols or onto surfaces may differ markedly in absolute quantity of infectious virus deposited. Here and in other studies, we use large initial quantities not because these are necessarily a realistic stand in for any or all depositions^11^, but rather because this enables maximally informative estimates of decay rates and half-lives. Since the decay process is approximately exponential, these rate estimates can be used for risk assessment for a wide range of deposition sizes.

### 8 Code and data

All code and data needed to reproduce the analyses described here is archived on Github (https://example.com) and Zenodo (https://example.com), and licensed for reuse, with appropriate attribution and citation.

## Notes

### Competing Interest Statement

The authors have declared no competing interest.

